# The O-glycosyltransferase SECRET AGENT Participates in Abscisic Acid-Induced Microtubule Remodeling and Stomatal Closure in Arabidopsis thaliana

**DOI:** 10.1101/2025.10.29.683829

**Authors:** Pengfang Sun, Yueran Wu, Pan Wang, Zixuan Wang, Miao Hu, Jing Li, Rong Yu

## Abstract

Secret Agent (SEC) encodes the major *O*-GlcNAc transferase in *Arabidopsis thaliana* and has been implicated in stress and signaling pathways, but its role in stomatal regulation remains unknown. In this study, we investigated the function of SEC in abscisic acid (ABA)-induced stomatal closure and drought responses. Loss of SEC function in the *sec*-*5* mutant delayed stomatal closure and increased water loss under drought, whereas *SEC* overexpression enhanced stomatal responsiveness and drought tolerance. Pharmacological assays revealed that *sec*-*5* was less sensitive to microtubule-stabilizing and -disrupting drugs during ABA-induced stomatal closing, consistent with a defect in cytoskeletal dynamics. Confocal imaging and quantitative analysis of microtubule rearrangements showed that SEC loss-of-function mutation retarded microtubule reorganization in response to ABA, with abnormal microtubule bundling and density. Mechanistically, SEC physically interacted with α-tubulin 4 (TUA4) in both yeast two-hybrid and bimolecular fluorescence complementation assays, suggesting tubulin as a potential substrate of SEC-modified *O*-GlcNAcylation. Together, these findings establish SEC as a key regulator of ABA-induced microtubule remodeling in guard cells, thereby linking *O*-GlcNAcylation to stomatal regulation and drought tolerance.

## INTRODUCTION

Stomata are microscopic pores formed by pairs of specialized epidermal cells referred to as guard cells. They are considered the primary gateways for gas exchange and water regulation between plants and their external environment, thereby playing a critical role in plant growth and survival. The evolution of stomata has been recognized as a pivotal milestone in the terrestrial adaptation of ancestral aquatic plants approximately 450 million years ago (Li *et al*., 2022). These structures facilitate CO_2_ uptake for photosynthesis and regulate water loss through transpiration, thus maintaining the plant’s internal water potential equilibrium (Torii, 2021). In agricultural crop species, stomatal regulation is vital for enhancing water use efficiency and resilience to abiotic treatment. At night, when photosynthesis ceases, stomata close to minimize unnecessary water loss. Drought stress, one of the most severe abiotic challenges confronting global agriculture, requires the activation of effective stomatal regulation strategies to enable plants to adapt and thrive under adverse environmental conditions (Gong *et al*., 2020).

Abscisic acid (ABA), a plant hormone, is a key regulator of plant responses to abiotic stress such as drought, high salinity, and low temperature. ABA is mainly synthesized in roots and subsequently transported to guard cells in leaves via ATP-binding cassette (ABC) transporter proteins, including AtABCG25 (ABC subfamily G 25), AtABCG26, and AtABCG40, or is locally synthesized through CLE25 (CLAVATA 3 (CLV3)/EMBRYO SURROUNDING REGION 25) signaling peptides within leaf tissues (Takahashi *et al*., 2018; Anfang & Shani, 2021). Following its synthesis and accumulation, ABA is perceived by pyrabactin resistance (PYR)/PYR1-like (PYL)/regulatory components of ABA receptor (RCAR) receptors in guard cells, thereby inhibiting group A protein phosphatases 2Cs (PP2Cs). This inhibition leads to the activation of sucrose non-fermenting 1 (SNF1)-related protein kinase 2 (SnRK2) family kinases, such as SnRK2.2, SnRK2.3, and SnRK2.6 (also referred to as OST1, OPEN STOMATA 1), which in turn activate a cascade of downstream response elements (Liu *et al*., 2022). These SnRK2 kinases phosphorylate various ion channels, including SLow Anion Channel 1 (SLAC1) and KAT1 (potassium channel protein), which regulate ion fluxes in guard cells, ultimately resulting in stomatal closure (Vahisalu *et al*., 2008; Sato *et al*., 2009).

In addition to ion channel regulation, the maintenance of the cellular cytoskeleton, particularly microtubules, within guard cells is critical for the regulation of stomatal dynamics. Microtubules are hollow, cylindrical protein fibers composed of α/β-tubulin heterodimers and are characterized by dynamic instability, manifested as rapid transitions between polymerization and depolymerization states. It has been demonstrated that microtubule depolymerization precedes stomatal closure during ABA-induced responses, and that the application of microtubule-stabilizing agents significantly inhibits this process (Dou *et al*., 2021). Consistently, genetic studies have identified several regulators of ABA-mediated microtubule reorganization. For instance, the *Arabidopsis* Really Interesting New Gene (RING) E3 ubiquitin ligase JAV1-ASSOCIATED UBIQUITIN LIGASE1 (JUL1) is required for ABA-induced microtubule depolymerization, stomatal closure, and drought tolerance (Yu *et al*., 2020), while the microtubule-associated protein MICROTUBULE-RELATED E3 LIGASE57(MREL57) mediates microtubule destabilization during ABA signaling (Dou *et al*., 2021). More recently, evidence has indicated that *O*-linked β-N-acetylglucosamine (*O*-GlcNAc) and *O*-linked fucose(*O*-fucose) modifications also participate in ABA signaling and microtubule reorganization in guard cells (Li *et al*., 2022). In parallel, large-scale proteomic analyses have revealed that the *O*-GlcNAc transferase SEC modifies hundreds of proteins in *Arabidopsis*, including tubulin and other regulatory components, thereby linking SEC activity to cytoskeletal regulation and stress signaling (Shrestha *et al*., 2024). Together, these findings suggest that microtubule organization and remodeling represent active processes that function as integral components of the ABA signaling pathway governing stomatal dynamics.

Post-translational modification, such as *O*-GlcNAc, has received growing attention as it regulates microtubule dynamics. This dynamic and reversible modification is primarily added to the hydroxyl groups of serine or threonine residues and is implicated in the regulation of signaling pathways such as mitogen-activated protein kinase (MAPK) and nuclear factor κB (NFκB) in animals (Mukherjee *et al*., 2007; Diallo *et al*., 2019). In plants, *O*-GlcNAc is catalyzed by the enzymes SECRET AGENT (SEC) and SPINDLY (SPY), both of which contain tetratricopeptide repeat (TPR) domains and *O*-GlcNAc transferase (OGT) catalytic domains, and SEC is notably conserved across both plant and animal species (Delporte *et al*., 2014). Uridine Diphosphate-N-acetylglucosamine (UDP-GlcNAc) is utilized by SEC as a donor substrate to catalyze *O*-GlcNAc modification of nuclear and cytoplasmic proteins, thereby functioning as a key regulator of plant development and hormone signaling (Olszewski *et al*., 2010; Mutanwad & Lucyshyn, 2022). And SPY is recently shown to be an *O*-fucosyltransferase (Zentella *et al*., 2017; Sun, 2021; Zentella *et al*., 2023).

In 2017, Xu et al. carried out a proteomic study to identify 262 *O*-GlcNAc-modified proteins in *Arabidopsis*, which encompassed functional categories such as transcriptional regulation, signal transduction, and tubulin family proteins (Xu *et al*., 2017). Studies in animal cells have shown that *O*-GlcNAc modification of α-tubulin can inhibit microtubule polymerization, thereby negatively regulating microtubule assembly (Ji *et al*., 2011).

Conversely, other research suggests that this modification may promote microtubule depolymerization, contributing to ciliary shortening (Tian & Qin, 2019). More recently, Liu et al. demonstrated that the *O*-fucosyltransferase SPY interacts with tubulin and regulates ABA-induced microtubule reorganization and stomatal closure in *Arabidopsis*, highlighting the importance of sugar modifications in microtubule-associated stomatal regulation (Liu *et al*., 2025). However, whether SEC regulates stomatal movement via the microtubule cytoskeleton remains unresolved and is an important question.

In this work, we demonstrate that SEC is required for ABA-induced stomatal closure and drought tolerance in *Arabidopsis* by regulating microtubule remodeling in guard cells. The *sec*-*5* mutant displayed reduced drought resistance, with accelerated water loss and delayed ABA-induced stomatal closure, while overexpression of SEC restored and enhanced these responses. Microtubule drug treatment revealed that *sec*-*5* was less sensitive to microtubule-stabilizing or -depolymerizing agents, and confocal imaging showed impaired microtubule depolymerization and bundling in guard cells. Furthermore, we show that SEC directly interacts with TUA4, suggesting tubulin as a potential *O*-GlcNAcylation target. Together, these findings establish SEC as a central regulator linking ABA signaling to microtubule dynamics, providing mechanistic insight into stomatal regulation and plant adaptation to drought stress.

## MATERIALS AND METHODS

### Plant Materials

*Arabidopsis thaliana* (Columbia-0, Col-0) was used as the wild-type (WT) ecotype, and all plant materials were maintained in the Col-0 background. The T-DNA insertion mutants with loss-of-function alleles of *SEC*, namely *sec*-*4* (Salk_106339) and *sec*-*5* (Salk_034290), were obtained from Dr. Fang Bao (Supplemental Fig. S1). The *SEC* overexpression lines OE#1 (*35Spro*:*SEC*#1) and OE#2 (*35Spro*:*SEC*#2) were a generous gift from Dr. Liangyu Liu. The *pTUB6::VisGreen*-*TUB6* construct was also kindly provided by Dr. Zhaosheng Kong. The *sec*-*5* (*pTUB6::VisGreen*-*TUB6*) line was generated in this study by crossing *pTUB6::VisGreen*-*TUB6* with the *sec*-*5* mutant. All mutant lines were confirmed to be homozygous by three-primer PCR genotyping (Supplemental Fig. S1). The primers used for genotyping are listed in Supplemental Tables S1–S3.

### Plasmid Construction

For the yeast two-hybrid (Y2H) assays, the coding sequence of *TUA4* was amplified using primers containing *EcoR*I and *BamH*I restriction sites and cloned into the *pGADT7* to generate the AD-TUA4 plasmid. The coding sequence of *SEC* was amplified using primers containing *BamH*I and *Pst*I restriction sites and cloned into the *pGBKT7* to generate the BD-SEC plasmid. The primers used for amplification are listed in Supplemental Tables S4.

For bimolecular fluorescence complementation (BiFC) assay, the coding sequence of *SEC* was amplified by PCR and inserted into *pCAMBIA*-*1300*-*nYFP* vector between *Xbal*I and *BamH*I sites. The coding sequence of *TUA4* was amplified by PCR and inserted into *pCAMBIA*-*1300*-*cYFP* vector between *Xbal*I and *BamH*I sites. The primers used for amplification are listed in Supplemental Tables S5.

### Reagents

The MES buffer consisted of 10 mmol L^-1^ MES, 100 μmol L^-1^ CaCl_2_, 50 mmol L^-1^ KCl, and 10 mmol L^-1^ KOH, adjusted to pH 6.1 with KOH. Abscisic acid (ABA) was diluted to 10 μmol L^-1^ in the epidermal strip buffer immediately before use. The microtubule-stabilizing agent Taxol and the microtubule-depolymerizing agent Oryzalin were each prepared at a concentration of 30 μmol L^-1^ and 10 μmol L^-1^ in the epidermal strip buffer prior to application.

### Plant Growth Conditions

*Arabidopsis* seeds were surface-sterilized sequentially in 70% ethanol and 25% sodium hypochlorite, rinsed thoroughly with sterile distilled water, and sown on 1/2 Murashige and Skoog (MS) medium. The seeds were vernalized at 4°C for 3 days and then transferred to a growth chamber under controlled conditions (light intensity: 120 μmol m^-2^ s^-1^; 16 h light/22°C, 8 h dark/18°C; relative humidity: 75%). After 10 days, seedlings were transplanted into soil and grown for approximately 3 weeks before being used in experiments.

### Drought Treatment Assay

For the drought treatment, 10-day-old seedlings of both WT and mutants were transplanted into moist soil and grown for two weeks under controlled conditions (light intensity: 120 μmol m^-2^ s^-1^; 16 h light at 22°C/8 h dark at 18°C; relative humidity: 75%). Irrigation was then withheld for approximately two weeks to impose drought stress. Plants were subsequently rewatered for three days, after which photographs were taken. Survival rates were calculated based on the number of plants showing leaf regreening and expansion. Each experiment was repeated at least three times.

### Detached Leaf Water Loss Assay

For water loss measurements, fully expanded rosette leaves from 3-week-old WT, mutants, and overexpression plants were excised from similar positions and at the same developmental stages. Detached leaves were placed on a laboratory bench at room temperature, and fresh weight was recorded every 30 min for up to 180 min. Water loss was expressed as a percentage of the initial fresh weight. Each experiment included 12 leaves per genotype and was repeated at least three times.

### Leaf Temperature Measurement

Thermal imaging was used to monitor leaf temperature as described (Wang *et al*., 2023; Zhong *et al*., 2024). Images of 3-week-old WT, mutants, and overexpression plants were acquired using an infrared thermal imaging camera (VarioCAM HD head 880), and false-colour infrared images were processed by IRBIS 3 Professional software. Measurements were conducted under identical ambient conditions in three independent experiments.

### Stomatal Aperture Measurement

All stomatal bioassays were performed as described, with some modifications (Dou *et al*., 2021; Wang *et al*., 2023). Rosette leaves of similar size and developmental stage were collected from 3-week-old plants and incubated in MES buffer under constant light (300 μmol m^-2^ s^-1^) for 120 min to promote stomatal opening. Leaves were then transferred to buffer containing the ABA (0 and 10 μM) with or without microtubule-specific drugs (30 μM Taxol or 10 μM Oryzalin) for an additional 120 min. Stomata were imaged using a ZEISS M200 upright fluorescence microscope, and pore widths were quantified from micrographs using ImageJ. For each genotype and treatment, at least 60 stomata were measured per biological replicate, with a minimum of three independent biological replicates.

### Yeast Two**-**Hybrid Assay

The full-length *SEC* coding sequence was cloned in-frame into the vector *pGBKT7* to generate BD–SEC, and the full-length *TUA4* coding sequence was cloned in-frame into the vector *pGADT7* to generate AD–TUA4. The constructs were co-transformed into *Saccharomyces cerevisiae* strain Y2HGold following the manufacturer’s instructions. Transformants were selected on synthetic dropout (SD) medium lacking leucine and tryptophan (SD/−Leu/−Trp) and then spotted onto SD medium lacking leucine, tryptophan, and histidine (SD/−Leu/−Trp/−His) to assess interaction via *HIS3* reporter activation. Plates were incubated at 28-30°C for 2-3 days before growth. Empty-vector combinations and single-plasmid transformations served as negative controls. All assays were performed in three independent experiments. The primers used for the yeast two-hybrid assay are listed in Supplemental Tables S4.

### Bimolecular Fluorescence Complementation Assay

The coding sequence of *SEC* was cloned into the *pCAMBIA*-*1300*-*nYFP* vector to produce SEC–nYFP, and the coding sequence of *TUA4* was cloned into the *pCAMBIA*-*1300*-*cYFP* vector to produce TUA4–cYFP. The plasmids were introduced into *Agrobacterium tumefaciens* GV3101 and injected into *Nicotiana benthamiana* leaves. GUS–nYFP and GUS– cYFP constructs served as negative controls. Plants were incubated in darkness for 12 h and then transferred to normal light conditions for 48 h. YFP fluorescence was detected using a fluorescence microscope. All assays were performed in three independent experiments. The primers are shown in Supplemental Tables S5.

### Quantitative Analysis of Microtubules Dynamics in Guard Cells

Leaves from 3-week-old *pTUB6::VisGreen–TUB6* transgenic *Arabidopsis* (WT and *sec*-*5*) were incubated in MES buffer under constant light (300 μmol m^-2^ s^-1^) for 120 min, then transferred to buffer containing the ABA (0 and 10 μM) for an additional 120 min. Abaxial epidermal strips were mounted and imaged on a Zeiss LSM 780 laser-scanning confocal microscope using a 63× oil-immersion objective (GFP excitation at 488 nm). Imaging parameters (laser power, detector gain/offset, pinhole, pixel size, and bit depth) were kept constant across treatments.

Quantification of microtubules dynamics in guard cells was performed as previously described (Higaki *et al*., 2010; Eisinger *et al*., 2012; Li *et al*., 2014; Ma *et al*., 2016; Dou *et al*., 2021; Wang *et al*., 2023; Zhong *et al*., 2024). The peak intensity of bundles in Type I microtubules was determined by measuring the continuous fluorescence intensity along the middle of the longitudinal direction of the guard cells (Fig. 6a), following the method of Eisinger *et al*. (2012). The number of microtubule filaments was quantified as the number of peaks with fluorescence intensity values greater than 50 (Li *et al*., 2014). To quantify microtubule density in guard cells, maximum-intensity projections were generated from serial confocal optical sections of *pTUB6::VisGreen*-*TUB6* transgenic lines. The projected images were binarized by thresholding and skeletonized using the ImageJ command “Process— Binary—Skeletonize.” The number of pixels corresponding to the skeletonized microtubules (*N_MT_*) and the total cell area (*N_cell_*) were measured in ImageJ. Microtubule density (100%) was calculated as 100 × *N_MT_* / *N_cell_*, representing the occupancy of Green fluorescence signals within guard cells (Higaki *et al*., 2010; Wang *et al*., 2023). To quantify the extent of microtubule bundling in guard cells, skewness of the intensity distribution of the microtubule pixels was measured as previously described (Higaki *et al*., 2010; Ma *et al*., 2016; Zhong *et al*., 2024).

Each parameter was quantified from at least 80 guard cells per genotype or treatment, and all analyses were repeated in three independent biological replicates.

### Statistical Analysis

All results are presented as means ± SD. Statistical analyses were performed using GraphPad Prism 10.0 software. Depending on the data distribution and experimental design, independent-samples two-tailed *t*-tests or one-way ANOVA were applied. Assumptions of normality and homogeneity of variances were assessed prior to analysis. For two-tailed *t*-tests, statistical significance was defined as \**P*<0.05, and high significance as \*\**P*<0.01. For one-way ANOVA, Different letters represent significant differences at *P*<0.05.

## RESULTS

### The *sec* Mutant Suppresses Drought Tolerance

To assess drought tolerance, 10-day-old *Arabidopsis* seedlings were planted into moist soil and grown for two weeks under controlled conditions. Irrigation was then withheld for approximately two weeks to impose drought stress. During this period, *sec*-*5* mutants displayed more severe and widespread phenotypic damage than other genotypes, such as extensive leaf desiccation, wilting, purple-brown discoloration, and eventual necrosis (Fig. 1a). Plants were then rewatered and survival rates were scored after 3 d of rewatering. As shown in Figure 1b, only 16.9% of the *sec*-*5* mutants were able to recover and resumed growth after re-irrigation, whereas much higher survival rates were observed in wild type (56.9%) and *sec*-*4* (∼47.8%). Statistical analysis revealed that there was no significant difference in survival rates between wild-type and *sec*-*4* mutant, but it was markedly reduced in *sec*-*5* seedlings, indicating that *sec*-*5* is more susceptible to water deficit than wild-type *Arabidopsis* plants. These results are consistent with the different *SEC* expression level in *sec*- *4* and *sec*-*5* by RT-qPCR test, showing that the expression of *SEC* in *sec*-*4* was similar to that in wild-type plants, while dramatically declined in the *sec*-*5* mutant (Supplemental Fig. S1c). These expression patterns about *sec*-*4* and *sec*-*5* have also been reported previously (Xing *et al*., 2018).

**Fig. 1.**
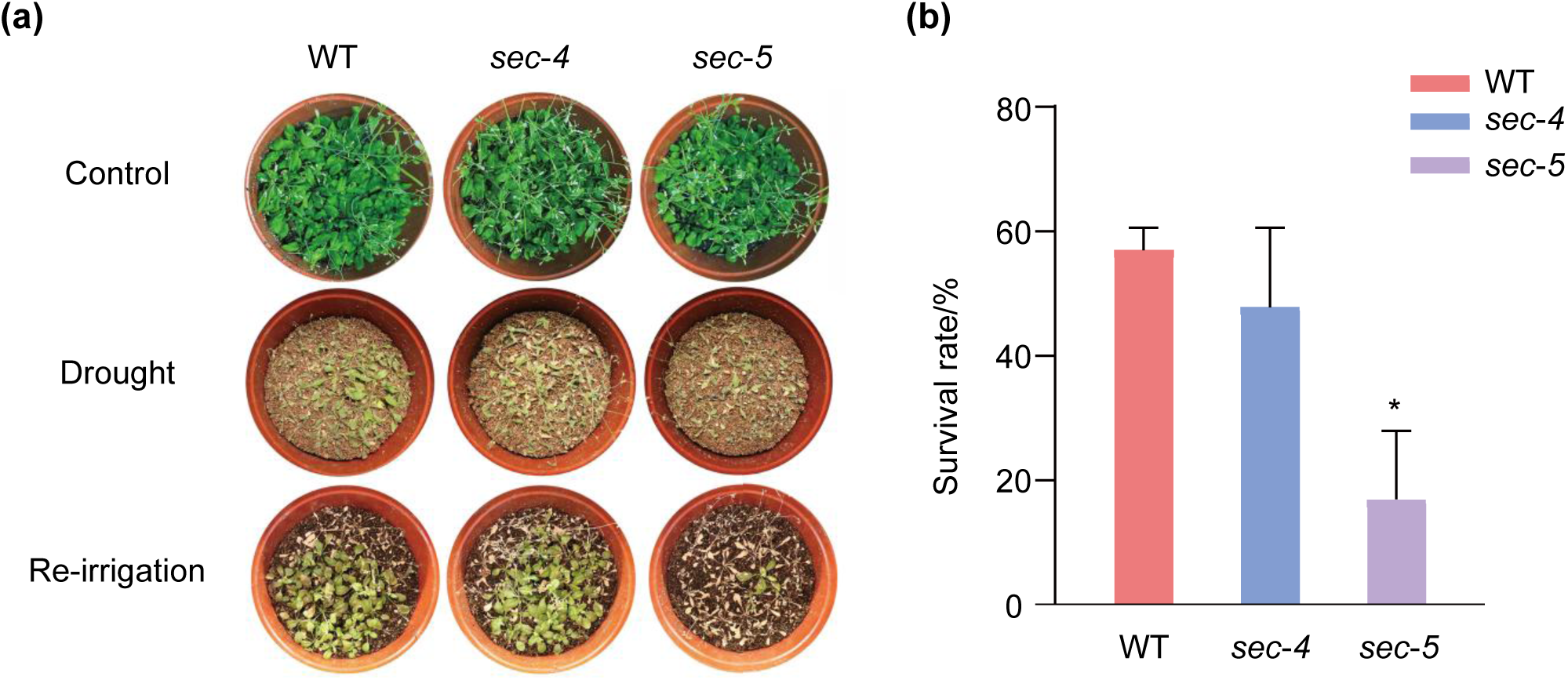
Phenotypic analysis of *sec* mutants in drought stress response. (a) Drought phenotypes of wild-type (WT), *sec*-*4*, *sec*-*5* seedlings grown in soil. 10-day-old seedlings were subjected to drought test by withholding water for 4 weeks before being photographed, and subsequently re-watered. (b) Quantification of the survival rate as shown in (a). Values represent means ± SD from three biological replicates. (Two-tailed *t* test, \**P*<0.05).

### Transpirational Water Loss is Faster in *sec***-***5* Compared with the Wild Type

Stomata, as the gatekeepers of uptake and defense signaling between terrestrial plants and the external atmosphere, the majority of plant water loss occurs through stomatal transpiration (Chua & Lau, 2024; Lawson & Leakey, 2024), evaluating leaf desiccation rate and infrared thermal imaging provides useful cues for intrinsic water use efficiency (Zheng *et al*., 2019; Dou *et al*., 2021; Wang *et al*., 2023; Li *et al*., 2024). To dissect the physiological role of SEC in stomatal movement precisely, two overexpression lines of *SEC* driven by the *35S* promoter (OE#1, OE#2) were recruited to perform transpiration-related assays, as well as the two T-DNA insertion mutants (*sec*-*4, sec*-*5*). RT-qPCR results confirmed considerably elevated expression levels of *SEC* in both overexpression lines compared to wild-type (Supplemental Fig.S1d). Leaves were collected from the 3-week-old plants of different genotypes, ensuring they were of the same developmental stages and leaf positioning.

Detached leaves were placed in a ventilated chamber to undergo natural dehydration, and fresh weights were recorded every 30 minutes. Notably, *sec*-*5* mutant leaves showed more pronounced wrinkling than those of wild-type plants, while *SEC*-OE lines exhibited relatively less wrinkling compared to both wild-type and *sec*-*5* mutant (Fig. 2a,b). Consistent with the survival rates in soil, no notable difference was observed in water loss between wild-type and *sec*-*4* leaves, which reached similar final rates of 46.8% and 42.5%, respectively. Conversely, leaves from *sec*-*5* mutants contrasted starkly with the faster water loss (53.2%). These results indicate that SEC is required for enhancing the ability of *Arabidopsis* to withstand drought stress.

**Fig. 2.**
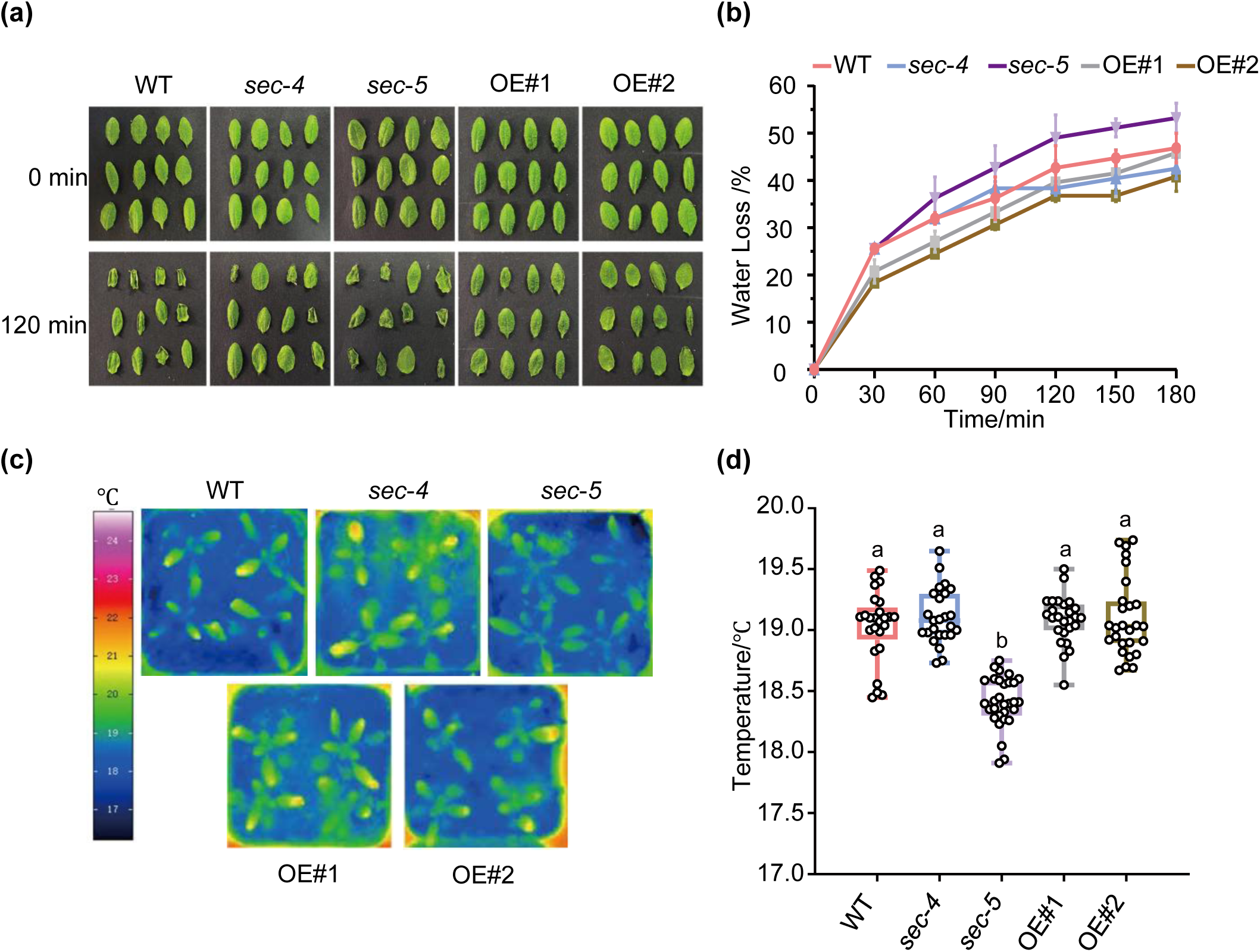
The *sec*-*5* mutant exhibits increased transpirational water loss. (a) The wilty phenotypes of detached rosette leaves of seedlings from WT, *sec*-*4*, *sec*-*5*, *SEC*-overexpression lines (OE#1, OE#2). The detached leaves were incubated on the foil on a laboratory bench (RH=35%). (b) Fresh weights of the detached leaves of WT, *sec*-*4*, *sec*-*5*, OE#1, OE#2 were measured designated time intervals. The experiment was repeated three times with independent treatments. Error bars indicate standard deviation. (c) Representative false-colour infrared images show that the leaf temperature of *sec*-*5* is lower than for the wild type. (d) Statistical analysis of leaf temperature of (c). A total of three independent experiments were performed and obtained similar results with a minimum of 25 leaves. Statistical significance was determined by one-way ANOVA. Different letters represent significant differences at *P*<0.05.

Given that increased transpiration rate typically results in decreased leaf surface temperature, we then surveyed leaf thermal profiles using an infrared thermal camera to determine the transpiration rate (Fig. 2c). As shown in Fig.2d, there was no significant difference in leaf temperature among wild-type, *sec*-*4*, and OE lines. However, leaf temperature of *sec*-*5* mutants was significantly lower compared to wild-type and other genotypes, in agreement with its higher transpiration rate and consequently reducing survival rate under drought conditions. In addition, considering the evidence about gene expression differences, soil drought phenotypes, water loss rates, and leaf temperatures between *sec*-*4* and *sec*-*5*, the *sec*-*5* mutant was then selected to be used for further functional analysis, since it carries an insertion in the second exon of the *SEC* genome (Supplemental Fig. S1a), resulting in a significant decrease in *SEC* transcript level and is considered the strongest allele (Xing *et al*., 2018; Shrestha *et al*., 2024).

### SEC Is Involved in ABA-Induced Stomatal Closure

Abscisic acid (ABA), one of the key phytohormones, plays a pivotal role in mediating plant responses to drought stress (Hsu *et al*., 2021; Kishor *et al*., 2022; Sharma *et al*., 2023). In *Arabidopsis thaliana*, *O*-GlcNAc glycosylation catalyzed by the *O*-GlcNAc transferase SEC has been implicated in multiple plant hormone signaling pathways, including gibberellin, cytokinin, auxin, and ABA (Xu *et al*., 2017). To investigate whether SEC-mediated modification also functions in ABA-induced stomatal closure, we employed *sec*-*5* mutant and overexpression lines (OE#1 and OE#2) for stomatal bioassay. As shown in Fig. 3a,b, after 120-minute ABA treatment, stomatal closure in wild-type plants showed approximately a 49% reduction, whereas *sec*-*5* mutants exhibited only about a 36% decrease. Although ABA signal led to some stomatal closure in *sec*-*5* leaves, their aperture still remained significantly wider than that of WT, indicating a diminished ABA responsiveness (Fig. 3b). Meanwhile, the OE#1 and OE#2 lines displayed similar stomatal movement to WT, with the 48% and 49% degrees of stomatal closure. Taken together, these findings suggest that SEC contributes positively to ABA-induced stomatal closure and that loss of its function impairs the normal stomatal response to ABA signaling.

**Fig. 3.**
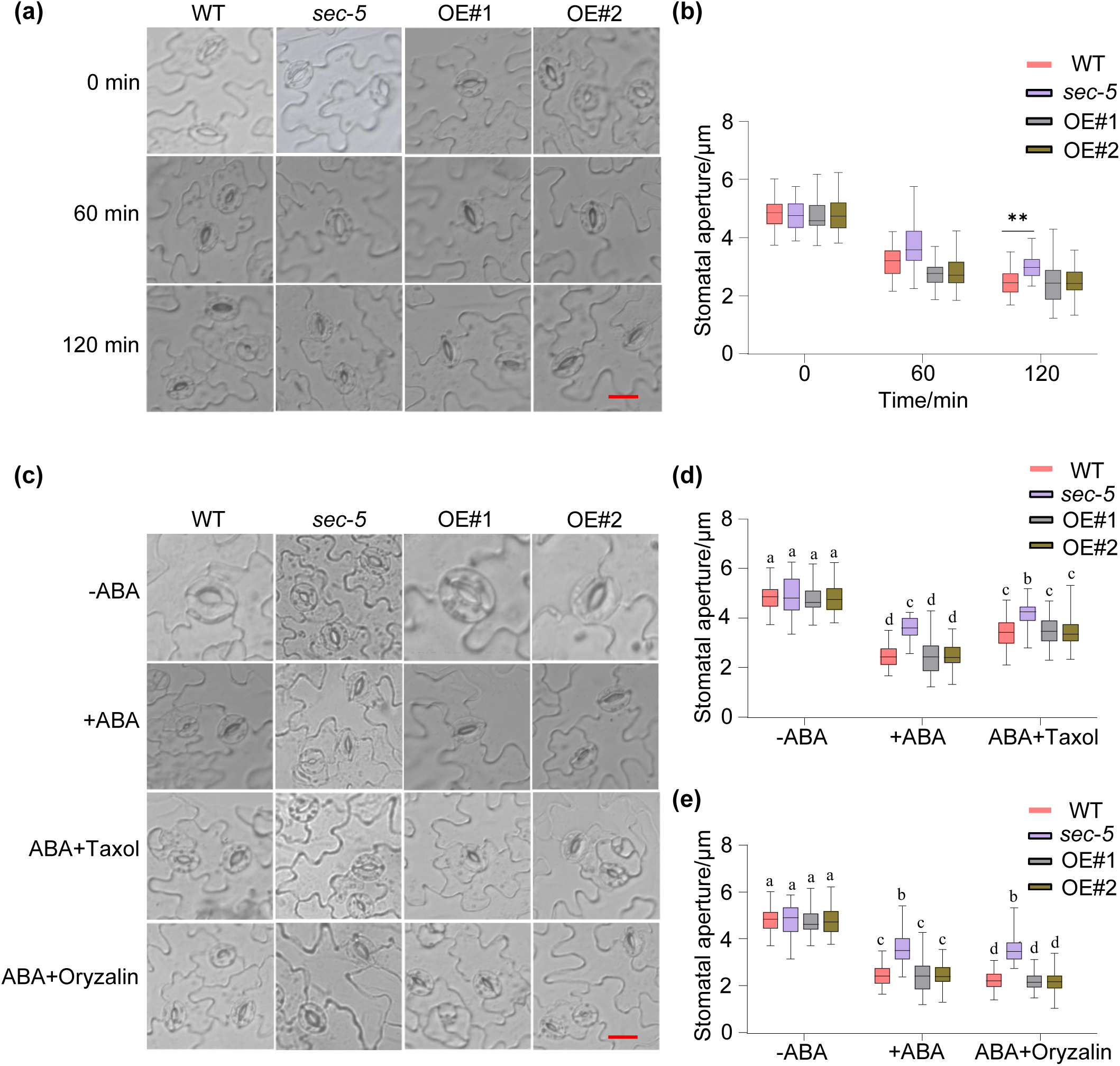
SEC regulates ABA-induced stomatal closure. (a) Representative images of stomatal aperture in wild type (WT), *sec*-*5* mutant and overexpression lines (OE#1 and OE#2) before and after treated with ABA treatment. A total of three independent experiments were performed and obtained similar results. Scale bars=20 μm. (b) Quantification of stomatal aperture size in wild type, *sec*-*5* mutant and overexpression lines of (a). A minimum of 60 stomatal pores were measured. (Two-tailed *t* test, *\*\*P*<0.01). (c) Representative images of stomatal aperture in WT, *sec*-*5* mutant and overexpression lines (OE#1 and OE#2) before and after treated with ABA and microtubule drugs (Taxol and Oryzalin). A total of three independent experiments were performed and obtained similar results. Scale bars=20 μm. (d, e) Quantification of stomatal aperture size in wild type, *sec*-*5* mutant and overexpression lines of (c). A minimum of 60 stomatal pores were measured. Statistical significance was determined by one-way ANOVA. Different letters represent significant differences at *P*<0.05.

It is well documented that microtubule dynamics is an indispensable component of ABA-induced stomatal closure (Yu *et al*., 2020; Dou *et al*., 2021; Wang *et al*., 2023). And a large-scale proteomics of SEC-mediated *O*-GlcNAcylation in *Arabidopsis thaliana* has unveiled numerous *O*-GlcNAc–modified proteins, including tubulin the subunit that formed microtubules, and ABA-signaling proteins (Xu *et al*., 2017; Shrestha *et al*., 2024; Aizezi *et al*., 2025). Given the potential relationship between SEC and tubulin, we reasoned that in guard cells, SEC might interact with tubulin by *O*-GlcNAcylation to regulate microtubule stability during ABA-induced stomatal movement. To address this possible cause, we performed microtubule-specific pharmacological stomatal bioassays with the microtubule-stabilizing drug Taxol and a depolymerizing agent Oryzalin.

We measured the stomatal aperture of all plants treated with ABA plus microtubule-specific drugs (Taxol or Oryzalin) (Fig. 3c). Co-treatment of Taxol attenuated ABA-induced stomatal closure in all genotypes; however, the inhibitory effect was significantly greater in wild type than in *sec*-*5* mutants, indicating a reduced sensitivity of *sec*-*5* to Taxol (Fig. 3d). In contrast, Oryzalin can depolymerize microtubule filaments and enhance ABA-triggered stomatal closure. When quantified, this effect was pronounced in wild-type plants but significantly diminished in *sec*-*5* (Fig. 3e), suggesting its impaired responsiveness to microtubule-disrupting conditions. In addition, SEC overexpression lines (OE#1 and OE#2) consistently exhibited a stronger stomatal phenotype than *sec*-*5* mutants throughout the assay. Together, these results indicate that loss of SEC might compromise the microtubule dynamics about assembly/disassembly in guard cells, thereby weakening ABA-driven stomatal closing in response to water deficit stress.

### SEC Interacts with TUA4

In *Arabidopsis thaliana,* SECRET AGENT (SEC) has previously been reported to *O*- GlcNAcylate hundreds of proteins involved in diverse cellular processes by proteomic analysis, among which is TUA4, an α-tubulin isoform. (Xu *et al*., 2017). To test whether SEC directly interacts with TUA4, we performed yeast two-hybrid (Y2H) assays. The GAL4 DNA-binding domain–SEC (BD-SEC) and GAL4 activation domain–TUA4 (AD-TUA4) constructs were co-transformed into yeast strain Y2HGold and plated on 2D double-dropout synthetic medium (-Leu/-Trp) and 3D triple-dropout selective medium (-Leu/-Trp/-His).

Yeast two-hybrid (Y2H) analysis revealed a strong interaction between the glycosyltransferase SEC and the tubulin isoform TUA4. As shown in Fig. 4a, only co-expression of the BD-SEC bait and AD-TUA4 prey constructs supported vigorous yeast growth on the selective medium (-Trp/-Leu/-His), as the interaction activated the *HIS3* reporter gene. This growth was absent in all negative controls, including yeast co-transformed with the two empty vectors *pGADT7* and *pGBKT7*, BD-SEC with *pGADT7*, or AD-TUA4 with *pGBKT7*, confirming the specificity of the SEC-TUA4 interaction. To examine the interaction of SEC with TUA4 in vivo, we conducted Bimolecular Fluorescence Complementation (BiFC) assays in *Nicotiana benthamiana* leaves. *SEC* fused to the N-terminal fragment of *YFP* (SEC-nYFP) and *TUA4* fused to the C-terminal fragment (TUA4-cYFP) plasmid vectors were generated, transformed into Agrobacterium, and transiently co-expressed into leaf epidermal cells. The yellow fluorescent protein (YFP) reconstitution signal was detectable only when the fusion constructs SEC-nYFP and TUA4-cYFP were both present (Fig. 4b). Together, these results revealed that SEC did physically interact with TUA4 both in yeast and in plant cells.

**Fig. 4.**
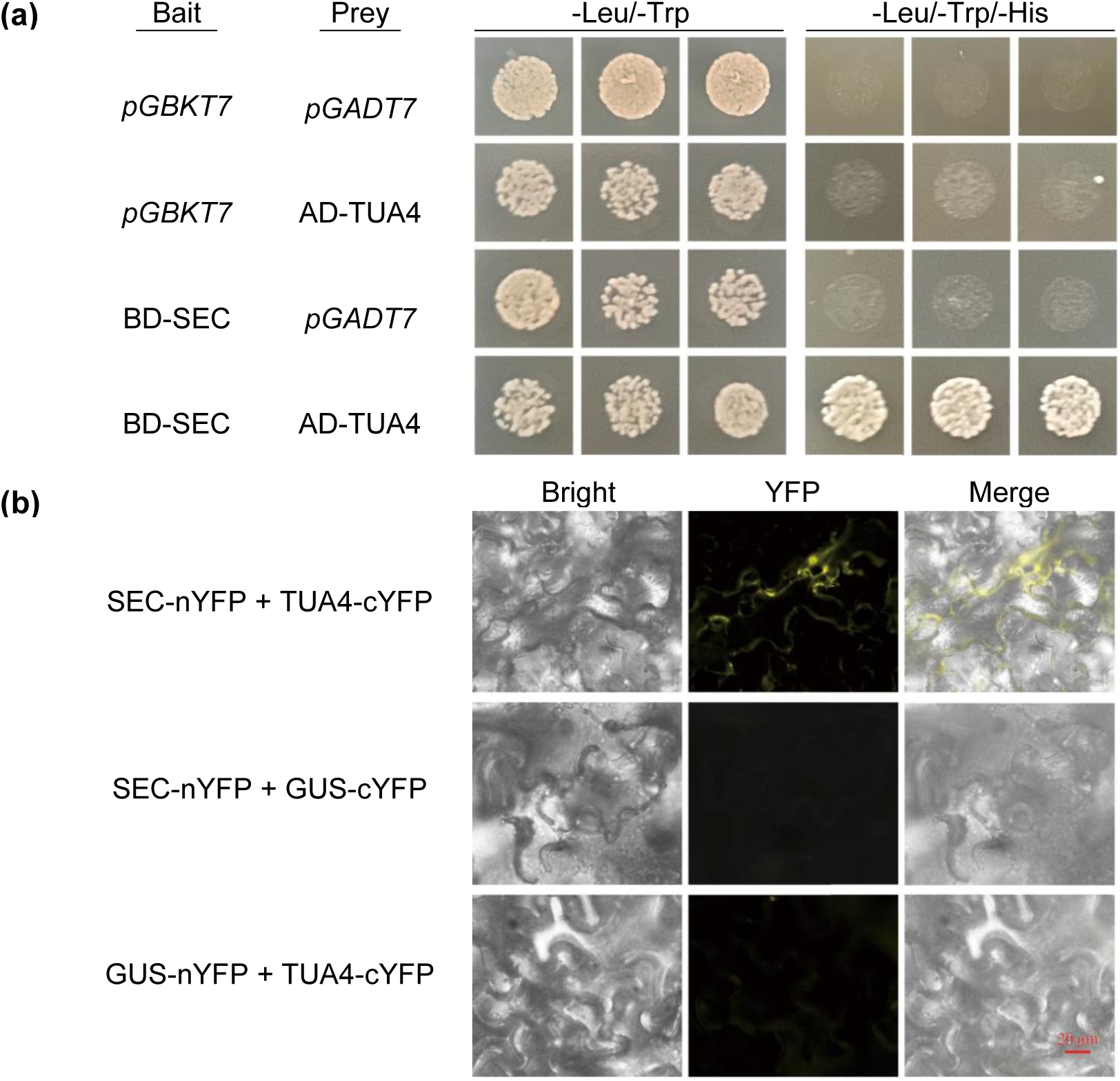
SEC interacts with Tubulin. (a) Y2H assay conducted to detect the interaction between SEC and TUA4. The full-length coding sequence (CDS) of *SEC* was cloned into the bait vector *pGBKT7* (BD). The full-length *TUA4* CDS was cloned into the prey vector *pGADT7* (AD). AD vectors expressing TUA4 was co-transformed with BD-SEC. SD/2, SD/-Leu/-Trp medium; SD/3, SD/-Leu/-Trp/-His medium. Yeast colonies were cultured on selective plates, and representative images from three independent yeast lines were captured after 3 d at 28°C. An empty vector was the negative control. (b) Analysis of the interaction of SEC with TUA4 using a bimolecular fluorescence complementation assay. Fusion protein expression vectors TUA4-cYFP and SEC-nYFP were constructed. GUS-cYFP and GUS-nYFP were used as negative control. The fluorescence of YFP was observed under fluorescence microscope. Scale bar=20 μm.

### The *sec***-***5* Mutant Exhibits Impaired ABA-Induced Microtubule Reorganization

Next, to assess the impact of SEC on ABA-induced microtubule remodeling in guard cells, we generated transgenic *Arabidopsis* harboring *pTUB6::VisGreen*-*TUB6* marker (Liu *et al*., 2019) in wild type (WT) and *sec*-*5* backgrounds. Since SEC participated in ABA-mediated stomatal closure (Fig.3) and directly interacted with the microtubule subunit-TUA4 (Fig.4), we reasoned that it might also be involved in ABA-mediated microtubule dynamics in guard cells. As expected, malfunction of SEC delayed typical microtubule disorganization in ABA signaling. Microtubule configurations in living guard cells were categorized into three types, as described by Fukuda *et al*. (1998): a highly organized radial state (Type I), a net-like state (Type II), and a disorganized fragmental state (Type III) (Fig. 5a)(Fukuda *et al*., 1998). After 120-min ABA treatment, a pronounced depolymerization of microtubules appeared in wild type guard cells, by a sharp decline in the proportion of cells with radial microtubule arrays (Type I) from 65.1% to 25.0%, a reduction reached 40.1 %. Concurrently, the percentage of cells containing disassembled or fragmented microtubules (Type III) rose markedly from 4.3% to 43.1%, an increase about 38.8%. By contrast, under the same conditions, the *sec*-*5* mutant displayed a delay in microtubule reorganization, in which radially arrayed microtubule filaments largely failed to depolymerize, only 10% guard cells of *sec*-*5* contained Type III arranged microtubules compared with 43.1% portion of wild-type. In *sec*-*5* stomata, filaments did not depolymerize to the degree seen in the WT (Fig.5b). These results indicate that the defect in SEC affects microtubule reorganization related to ABA-induced stomatal closure.

**Fig. 5.**
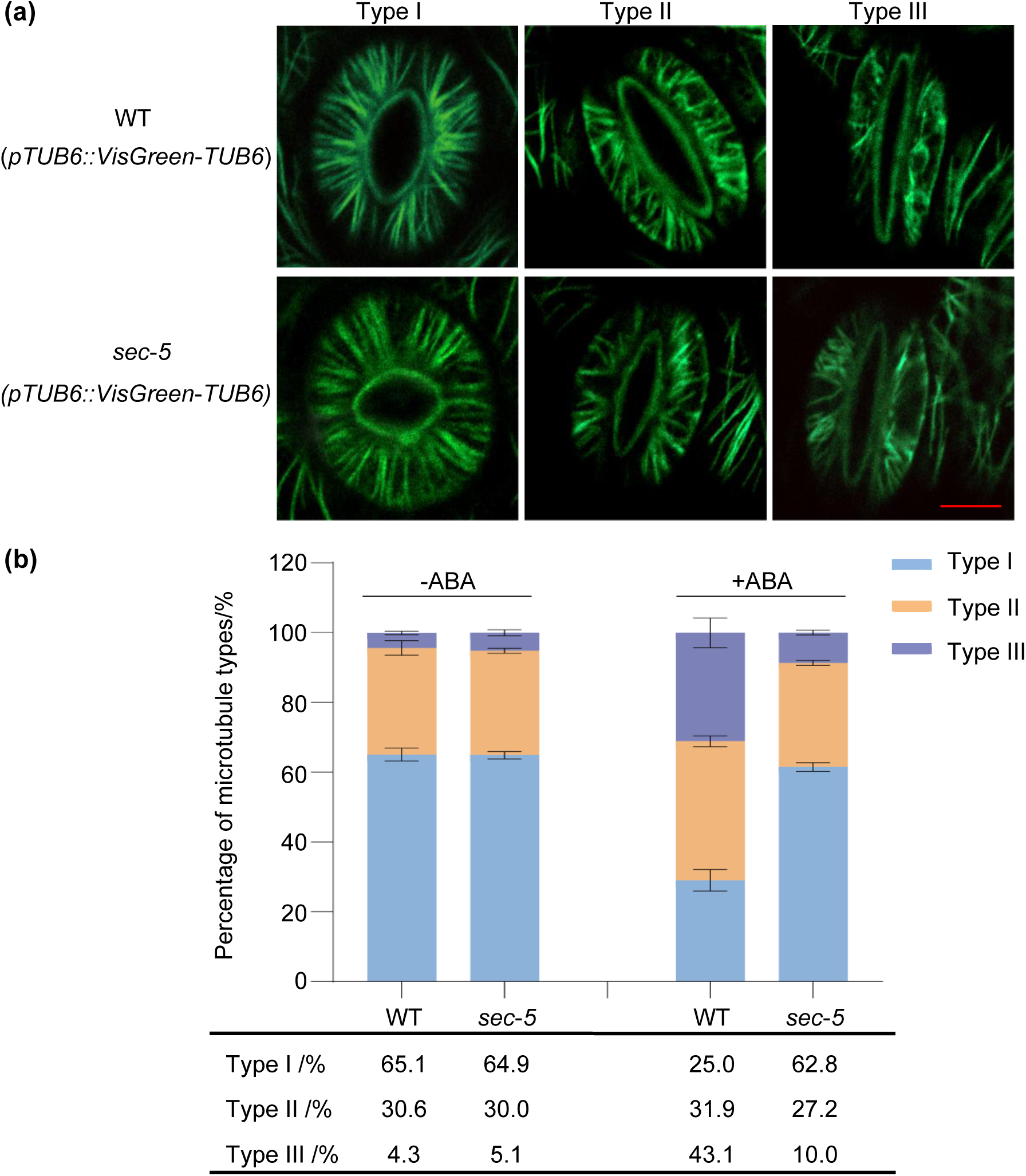
ABA-induced microtubule reorganization is suppressed in *sec* mutant. (a) Confocal images of microtubules in wild type and *sec*-*5* mutant guard cells during ABA-induced stomatal closure. Microtubules were classified into three groups: Type I (straight radial state), Type II (net-like state), and Type III (disassembled or fragmented state). Scale bar=5 μm. (b) proportions of different microtubule patterns in guard cells after ABA treatment. Values represent the means ±SD for three independent experiments, with at least 200 guard cells examined.

### Quantitative Analysis Reveals Increased Microtubule Number and Density in *sec***-***5* Guard Cells

To further characterize the differences in the configuration of microtubule filaments during ABA-triggered stomatal closure, we performed a quantitative analysis of several key parameters using ImageJ, and calculated the number of resolvable microtubules (Green fluorescence intensity > 50), MT density (Occupancy), and MT bundling (Skewness). These parameters have been used previously to quantify actin and microtubule changes in guard cells (Higaki *et al*., 2010; Eisinger *et al*., 2012; Li *et al*., 2014; Qian *et al*., 2019; Biel *et al*., 2020; Zou *et al*., 2021; Wang *et al*., 2023; Moser *et al*., 2024; Liu *et al*., 2025). As shown in Fig. 6a, filament number is defined as the number of fluorescent peaks along the mid-width line of the guard cell, as described by Li *et al*. (2014)(Li *et al*., 2014). It showed that wild type contained about 15 microtubule bundles per cell with a fluorescent intensity > 50 counted, whereas *sec*-*5* guard cells had a mean of 21 microtubule bundles per guard cell, which was significantly more abundant than WT (Fig. 6b), although the radial pattern of microtubules in wild-type looked similar to those in the *sec*-*5* mutant.

**Fig. 6.**
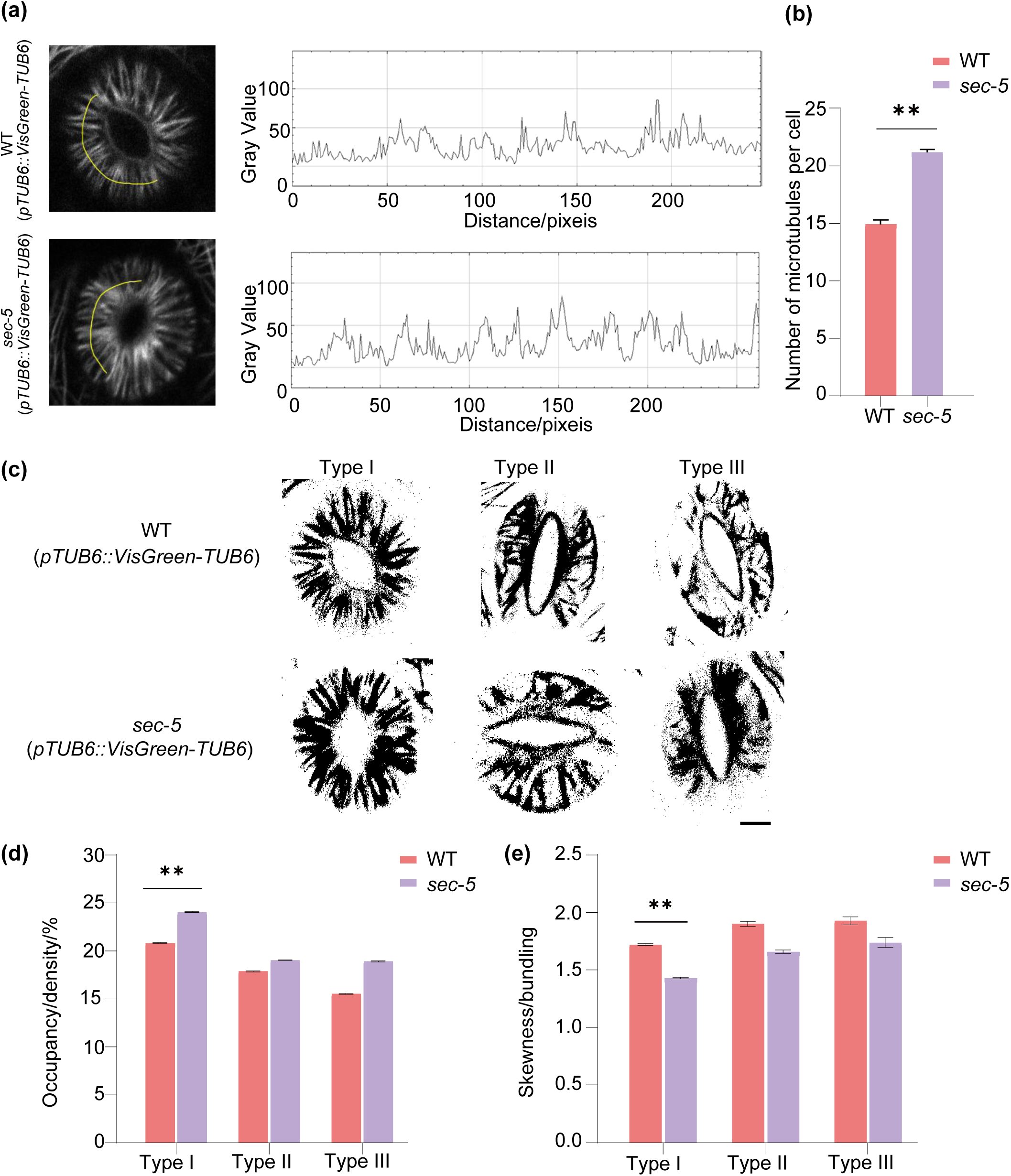
SEC regulates the dynamic rearrangement of microtubule in guard cells under ABA signaling. (a) Fluorescence intensity profiles of Green signals along a line in guard cells with radial microtubule orientation were measured in WT/*pTUB6::VisGreen-TUB6* and *sec-5*/*pTUB6::VisGreen-TUB6*. Scale bar=5 μm. (b) Quantification of microtubules with fluorescence intensity values greater than 50 (gray value) in (a). At least 60 stomatal pores were analyzed. The experiment was repeated three times with independent treatments (two-tailed t-test, \*\**P*< 0.01). (c) Representative images of guard cells after skeletonization using ImageJ. Scale bar=5 μm. (d) Statistical analysis of the occupancy of microtubules in guard cells. Three independent experiments were performed, yielding similar results, with at least 100 guard cells analyzed (two-tailed *t*-test, \*\**P*< 0.01). (e) Quantification of bundling (skewness) of microtubule filaments in WT and *sec-5* guard cells. Data presented are the means ±SD of three independent biological replicates. At least 100 guard cells were analyzed. (two-tailed *t*-test, \*\**P*< 0.01).

In addition, we evaluated the microtubule density by defining green fluorescent signal occupancy (displayed in %) using Image J software, to measure the average signal distribution within a guard cell. Analysis of skeletonized MT arrays (Fig. 6c) showed that MT density (occupancy) in the *sec*-*5* mutant was also significantly increased than in the wild type (Fig. 6d), consistent with the visually scored amount of microtubules (Fig. 6b). Moreover, the extent of microtubule bundling, as assessed via skewness analysis (Li *et al*., 2014; Zou *et al*., 2021; Zhang *et al*., 2024), was also calculated. In *sec*-*5* mutant, the skewness values exhibited a general reduction in microtubule bundling compared to the wild-type, with the most pronounced difference observed in radially arranged (Type I) microtubules (Fig. 6e). Thus, these analyses suggest that SEC are involved in modulating microtubule dynamics in guard cells, and a more dense and less bundled microtubule array is noted when SEC is absent in plants. Suppression of *SEC* probably hindered ABA-promoted microtubule depolymerization in guard cells, and consequently retarded stomatal closure in the *sec*-*5* mutant progeny.

## Discussion

With the currently changing climate and global warming, drought stress is becoming increasingly severe, which profoundly impacts plant growth and development, and restricts sustainable crop production. How to communicate in plants and enhance survival within fluctuating environments depends largely on the understanding of the synergistic effect of regulatory pathways, such as transcription factors, phytohormone, stomatal movement and so on (Chen & Ham, 2022; Cao *et al*., 2024; She *et al*., 2024; Zhang *et al*., 2024). In this work, we show that SEC plays a critical role in plant drought tolerance by mediating ABA-induced microtubule reorganization in guard cells. Loss of SEC function in the *sec*-*5* mutant impaired microtubule depolymerization, delayed stomatal closure, and reduced drought tolerance, whereas overexpression of *SEC* enhanced these responses. We further demonstrated that *sec*-*5* mutants exhibited reduced sensitivity to both microtubule-stabilizing and -destabilizing agents, consistent with a defect in microtubule dynamics. Moreover, we found that SEC directly interacts with the microtubule protein TUA4, suggesting that tubulin is a likely substrate of SEC-mediated *O*-GlcNAcylation. Together, these findings establish SEC as a key regulator linking ABA signaling to cytoskeletal remodeling, thereby offering mechanistic insights into stomatal regulation under drought stress. A working model of this regulatory pathway is shown in Fig. 7.

**Fig. 7.**
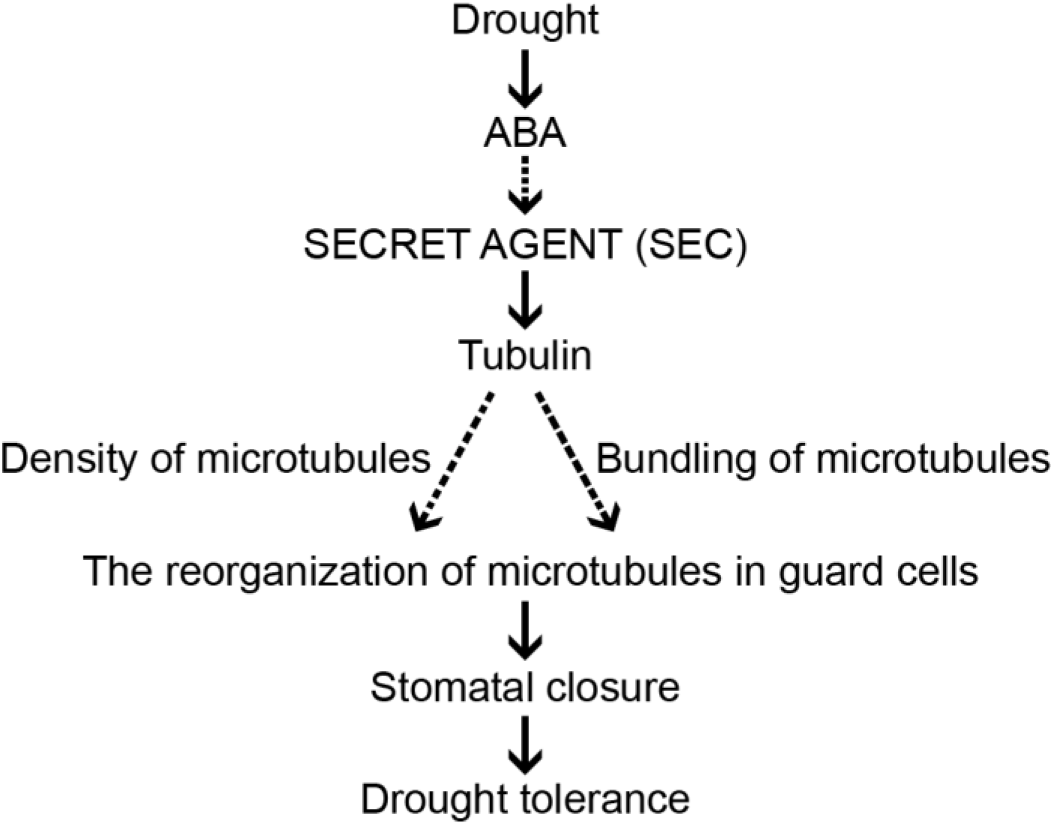
A model of the role of SEC in ABA-induced stomatal closure. Under drought stress, ABA signaling activates the glycosyltransferase SECRET AGENT (SEC), which modifies tubulin. SEC-dependent regulation affects both the density and bundling of microtubules in guard cells, thereby modulating their reorganization. This cytoskeletal remodeling promotes ABA-induced stomatal closure, leading to reduced water loss through transpiration and enhanced drought tolerance.

Research on *O*-GlcNAcylation has expanded beyond *Arabidopsis*, with growing evidence from crop species such as rice. For example, Li *et al*. reported that *O*-GlcNAcylation plays critical roles in rice growth and stress adaptation, suggesting that this modification is evolutionarily conserved across plant lineages (Li *et al*., 2023). Moreover, proteomic studies in *Arabidopsis* have identified hundreds of potential *O*-GlcNAc substrates, including signaling proteins, transcriptional regulators, and cytoskeletal components (Xu *et al*., 2017; Shrestha *et al*., 2024). However, only a few of these proteins have been functionally characterized in planta. As highlighted by Aizezi *et al*., most candidates remain unexplored, underscoring the gap between omics-level discovery and mechanistic understanding (Aizezi *et al*., 2025). Our work addresses part of this gap by validating α-tubulin as an SEC- interacting protein in guard cells and establishing its role in ABA-induced stomatal closure.

Our results further expand the current understanding of glycosylation in guard cell signaling. Previous work has shown that SPY, a fucosyltransferase, modifies tubulin and regulates ABA-induced microtubule reorganization during stomatal closure (Liu *et al*., 2025). In addition, proteomic studies identified α-tubulin as a potential *O*-GlcNAc-modified protein in *Arabidopsis* (Xu *et al*., 2017). While SPY-mediated *O*-fucosylation has been implicated in cytoskeletal regulation, our study demonstrates that SEC-mediated *O*-GlcNAcylation also participates in this process. These findings align with the broader concept that glycosylation functions as a critical regulatory layer in plant stress signaling (Bi *et al*., 2023). Importantly, our study highlights SEC as a previously underexplored regulator of tubulin function, thereby broadening the framework of sugar modification–mediated cytoskeletal regulation in plants.

Mechanistically, the impaired depolymerization observed in *sec*-*5* mutants suggests that SEC activity is required to destabilize microtubules during ABA signaling. This observation is consistent with studies in animal systems, where *O*-GlcNAc modification of α-tubulin has been reported to influence microtubule assembly and stability (Ji *et al*., 2011; Tian & Qin, 2019). Our results support the notion that SEC-dependent *O*-GlcNAcylation fine-tunes the balance between polymerization and depolymerization of guard cell microtubules, thereby enabling the dynamic cytoskeletal remodeling necessary for stomatal closure. This mechanism also explains the reduced responsiveness of *sec*-*5* mutants to both microtubule-stabilizing and -disrupting agents, as well as their impaired stomatal response to ABA.

Stomatal regulation is increasingly recognized as a central determinant of plant adaptive strategies under stress. In addition to their classical role in gas exchange and water loss, stomata integrate multiple environmental signals and act as decision-making hubs in stress responses (Tarkowski & Signorelli, 2025). While stomatal closure enhances drought tolerance and pathogen resistance, it also restricts CO_2_ uptake and volatile-mediated signaling, highlighting the physiological trade-offs plants face. Recent efforts in stomatal engineering have therefore shifted from altering stomatal density to improving stomatal kinetics, as faster opening and closing can enhance both water-use efficiency and carbon assimilation without compromising growth (Nguyen *et al*., 2023). In this context, our finding that SEC regulates microtubule reorganization during ABA-induced stomatal closure adds a mechanistic layer to understanding how post-translational modifications fine-tune stomatal dynamics. By linking *O*-GlcNAcylation to guard cell cytoskeletal remodeling, this work situates SEC within the broader framework of stomatal control that balances water conservation, carbon fixation, and stress defense.

The relationship between SEC and SPY represents another key frontier. SEC and SPY are the two canonical monosaccharide transferases in *Arabidopsis*, with extensive evidence indicating that they regulate growth and signaling in either synergistic or antagonistic ways (Mutanwad *et al*., 2021; Zentella *et al*., 2023). Our findings, together with those of Liu et al., suggest that SEC and SPY may also coordinately influence guard cell microtubule organization and stomatal dynamics (Liu *et al*., 2025). Future research should therefore examine whether these two enzymes exert cooperative or opposing functions in stomatal regulation. Given that SEC and SPY double mutants are lethal, such studies could leverage chemical inhibitors like SOFTI (SPY *O*-fucosyltransferase inhibitor), which has been developed as a tool to study *O*-fucosylation in plants (Aizezi *et al*., 2024). Combining SEC mutants with SOFTI treatment could mimic the double-mutant condition and reveal overlapping or distinct postembryonic functions of the two enzymes.

Despite these advances, our study has several limitations. Due to technical constraints, we were unable to directly detect *O*-GlcNAc modifications of TUA4 or other tubulin isoforms in planta, leaving the precise biochemical substrates of SEC unresolved. Future studies employing advanced glycoproteomic approaches will be critical to confirm whether tubulin is directly glycosylated by SEC. Additionally, while our work focused primarily on the microtubule cytoskeleton, SEC may also regulate other ABA-responsive proteins or actin filaments, which may further contribute to stomatal dynamics and drought responses. Dissecting these broader regulatory networks will require complementary biochemical and genetic approaches.

Taken together, our findings provide new insights into how SEC contributes to the regulation of guard cell function under drought stress. By linking ABA signaling to microtubule reorganization, SEC emerges as a crucial component of the stomatal closure machinery. This study not only underscores the importance of protein *O*-GlcNAcylation in cytoskeletal regulation but also introduces SEC as a novel player in plant stress physiology. The identification of SEC as a tubulin-interacting protein with a role in ABA-induced stomatal closure represents an important advance in understanding the post-translational control of microtubule dynamics. These findings highlight SEC as a promising target for genetic and biotechnological approaches to enhance crop resilience to drought stress.

## Acknowledgements

We thank Dr. Shanjin Huang (Tsinghua U.) and Dr. Tonglin Mao (China Agricultural U.) for helpful advice on the research design and comments on the article. We thank Dr. Zhaosheng Kong (Institute of Microbiology, The Chinese Academy of Sciences) for providing *Arabidopsis* seeds that expressing *pTUB6::VisGreen*-*TUB6*. We thank Dr. XingChen (Peking U.), Dr. Liangyu Liu and Dr. Fang Bao from Capital Normal University for providing experimental materials and for their guidance in the experiments.

## Author contributions

Pengfang Sun: investigation, writing. Yueran Wu: investigation. Pan Wang: investigation. Zixuan Wang: investigation. Miao Hu: investigation. Jing Li: conceptualization, resources, supervision, writing. Rong Yu: funding, conceptualization, data curation, supervision, project administration, writing.

## Data and resource availability

All relevant data can be found within the article.

## Supporting Information

Additional supporting information can be found online in the Supporting Information section.

---

**Supplemental Fig. S1.**
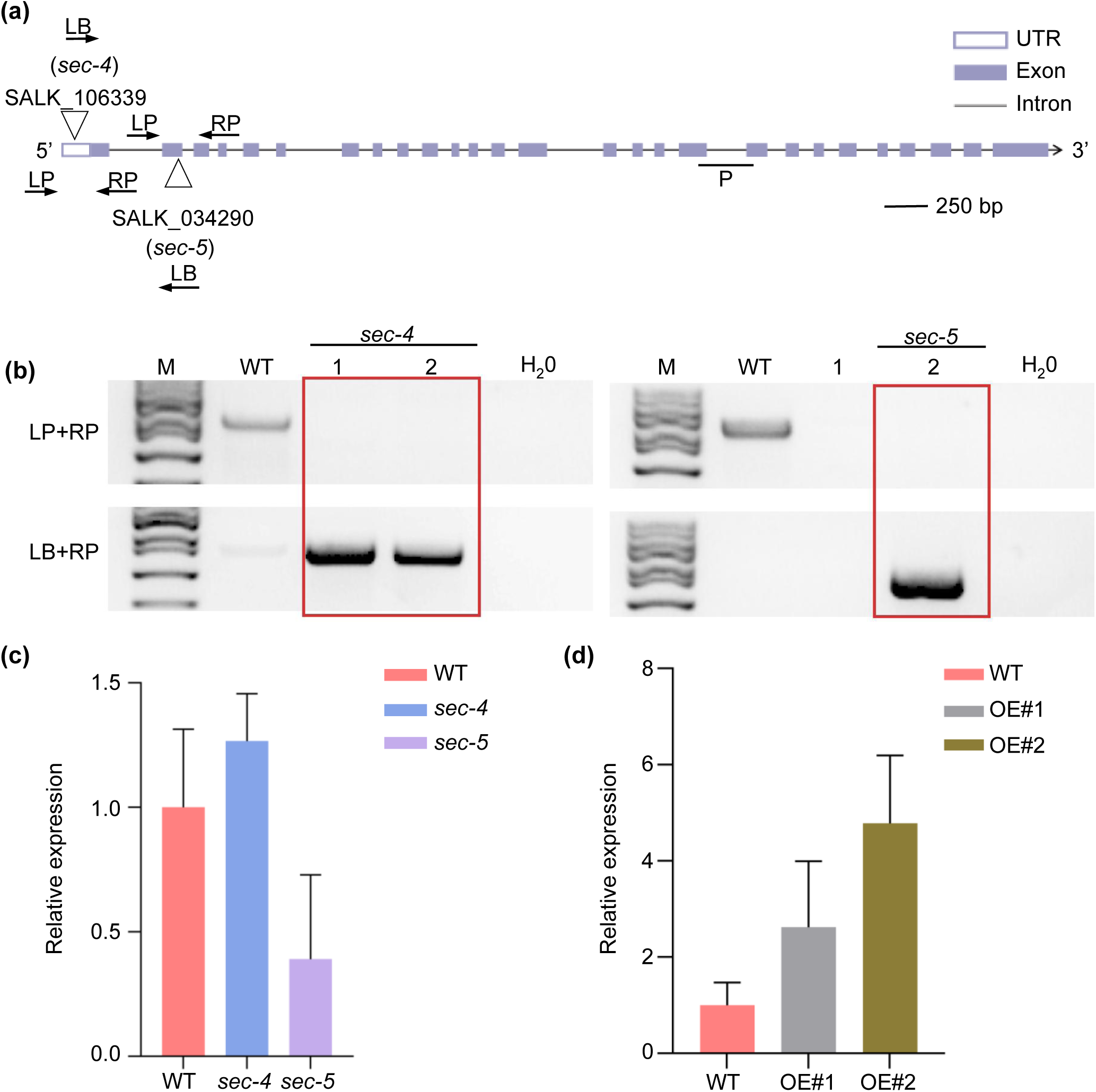
Identification of *sec* mutants. **(a)** Schematic genomic structure of the SEC locus. Exons are shown as purple boxes, introns are indicated as lines, UTR is shown as purple black box. T-DNA insertion positions are indicated by triangles. LB represents the left border primer of the T-DNA insertion. LP and RP represent the left and right genomic primers, respectively. P represents the SEC transcript detected by RT-qPCR. **(b)** Identification of *sec* homozygotes. (M: Marker. The molecular weight is 5000 bp. The lane 1 and 2 in the left panel are *sec-4* mutants. The lane 2 in the right panel is the *sec-5* mutant). **(c)** RT-qPCR analysis of SEC mRNA levels in *sec-4* and *sec-5* mutants. The expression level was normalized to that of *EF1-α*, a reference gene for RT-qPCR. Independent biological experiments were repeated three times. Data shown are means ± SD. **(d)** RT-qPCR analysis of *SEC* mRNA levels in *SEC*-overexpression (OE) lines. The expression level was normalized to that of *EF1-α*, a reference gene for RT-qPCR. Independent biological experiments were repeated three times. Data shown are means ± SD.

