## Supplemental Figure S1 for "The O-glycosyltransferase SECRET AGENT Participates in Abscisic Acid-Induced Microtubule Remodeling and Stomatal Closure in Arabidopsis thaliana"

**Supporting Information**

**Fig. S1. Identification of *sec* mutants.**

SALK_106339

Exon

Intron

UTR

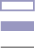

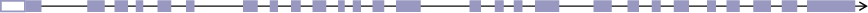

(*sec-4*)

SALK_034290

(*sec-5*)

LB

RP

LP

LB

RP

LP

5’

3’

250 bp

**(a)**

**(b)**

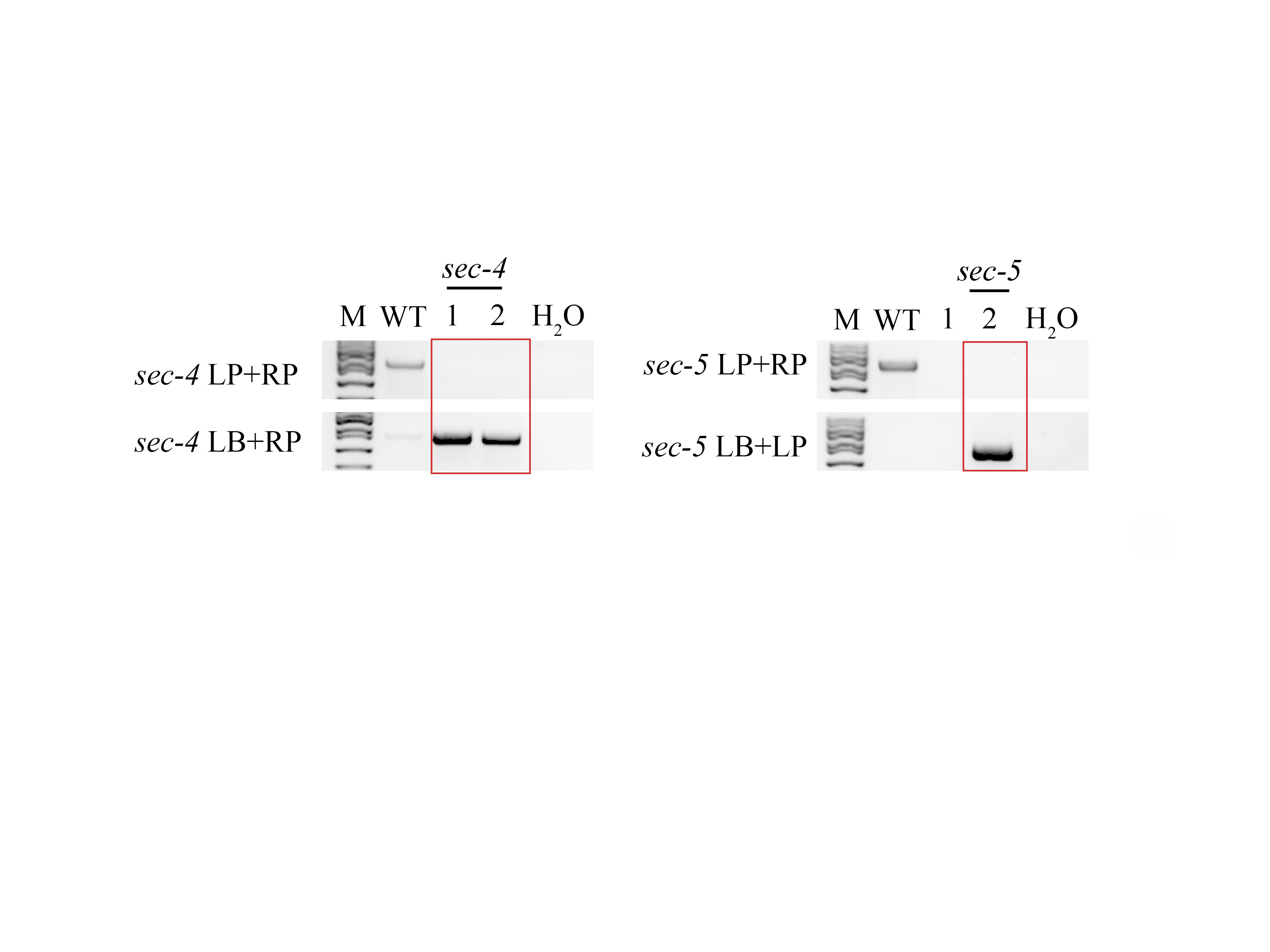

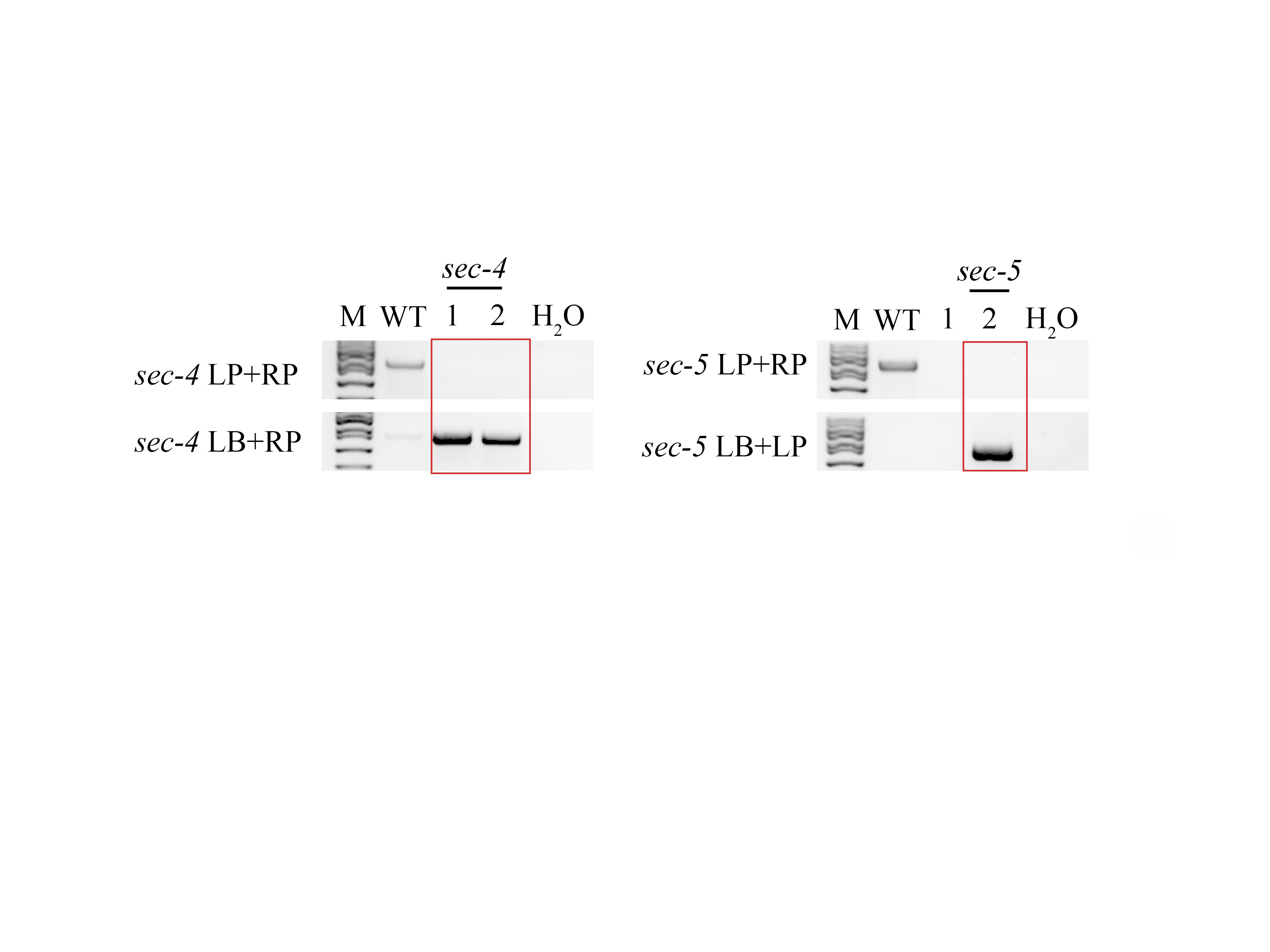

LP+RP

LB+RP

M

WT

1

2

H_2_0

M

WT

1

2

H_2_0

**(c)**

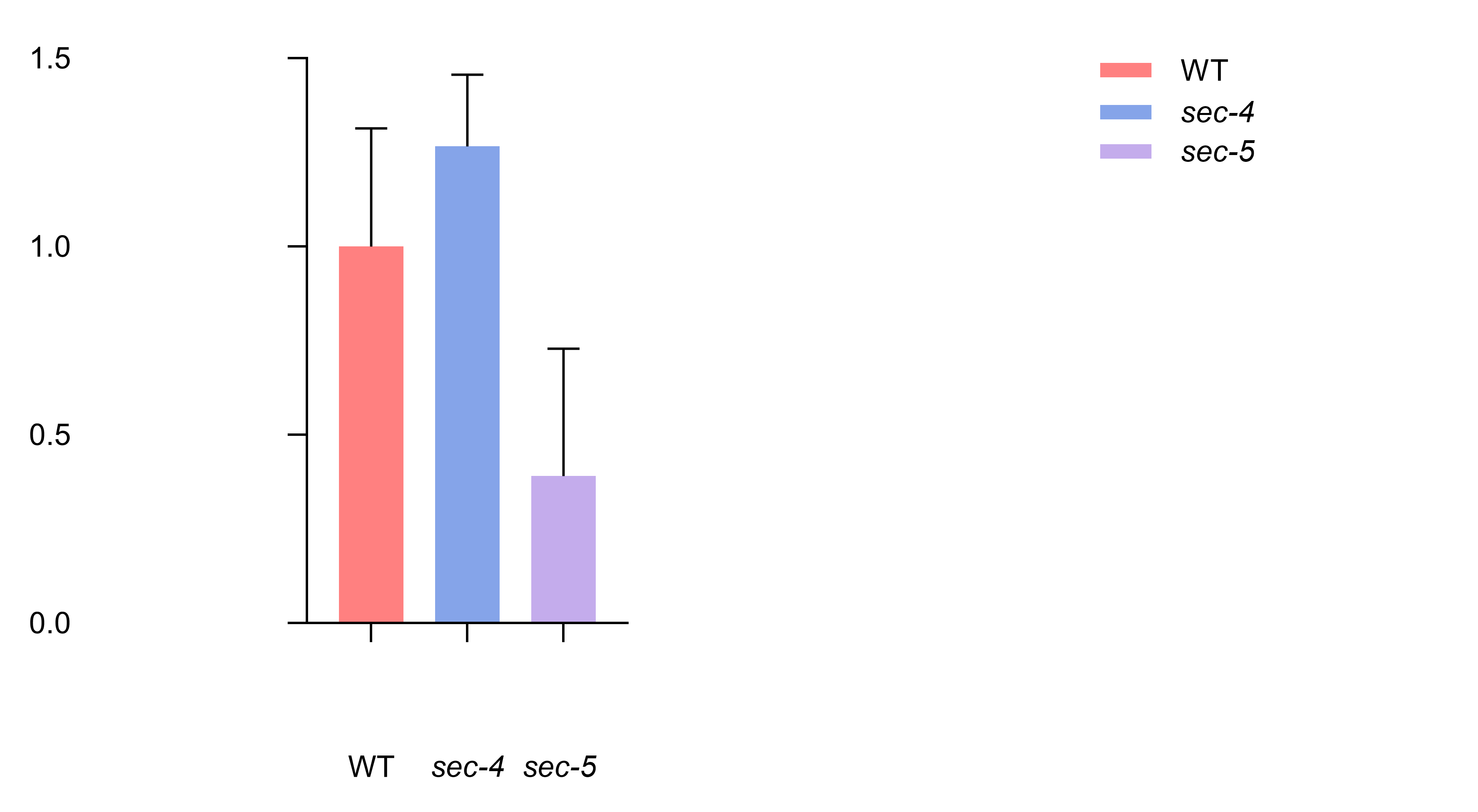

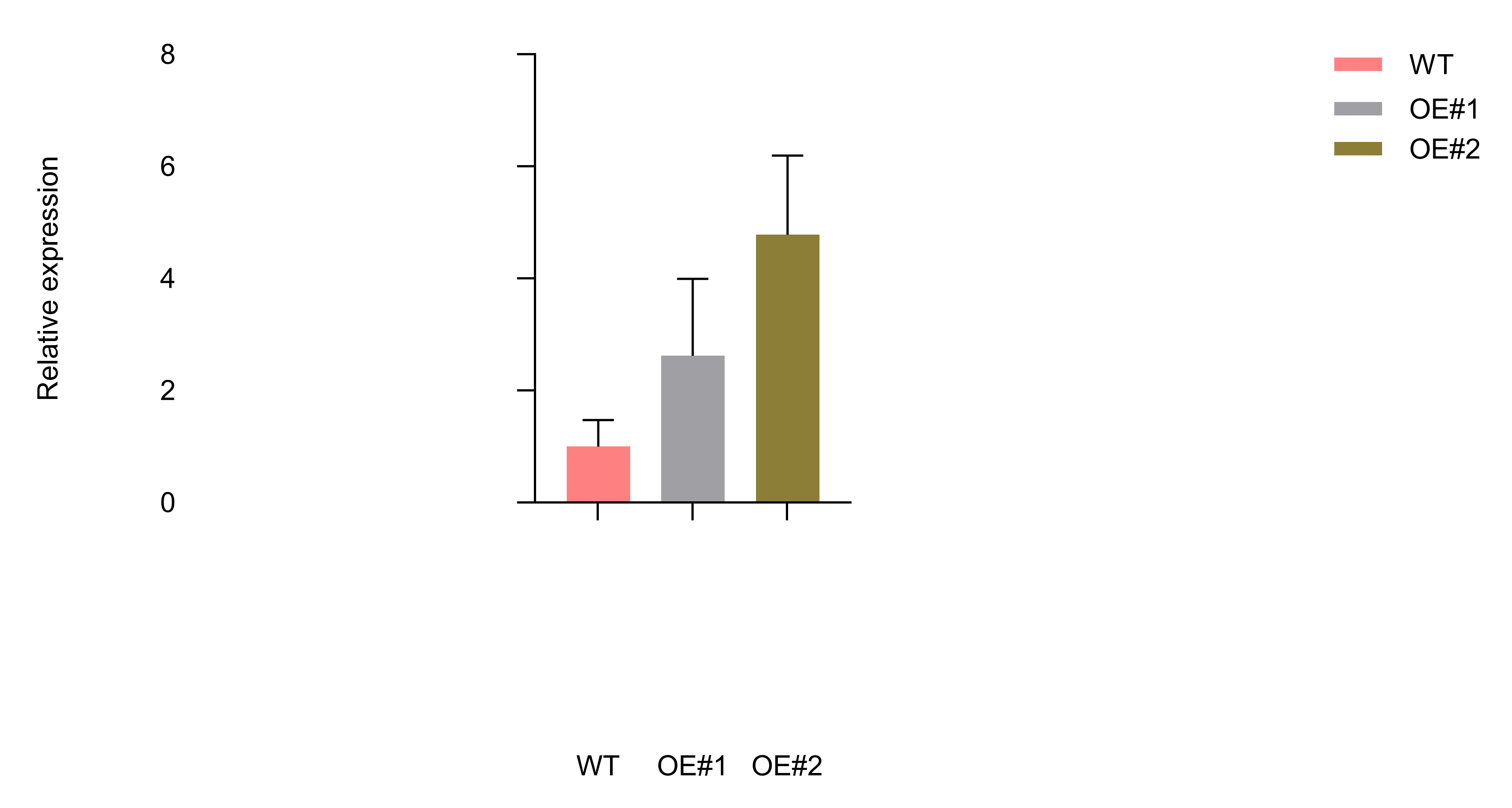

**(d)**

0.0

0.5

1.0

1.5

WT

*sec-4*

*sec-5*

Relative expression

Relative expression

0

2

4

6

8

WT

OE#1

OE#2

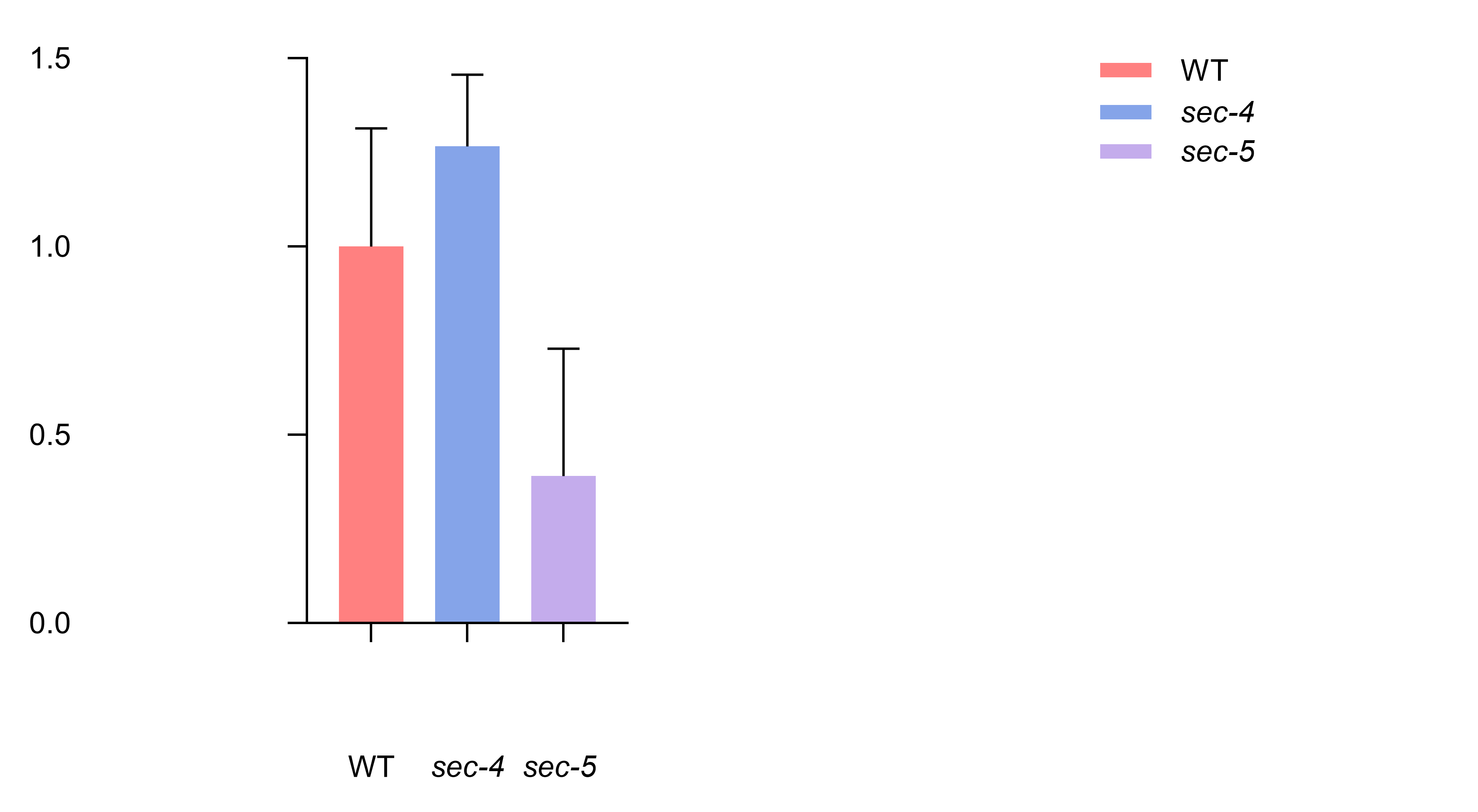

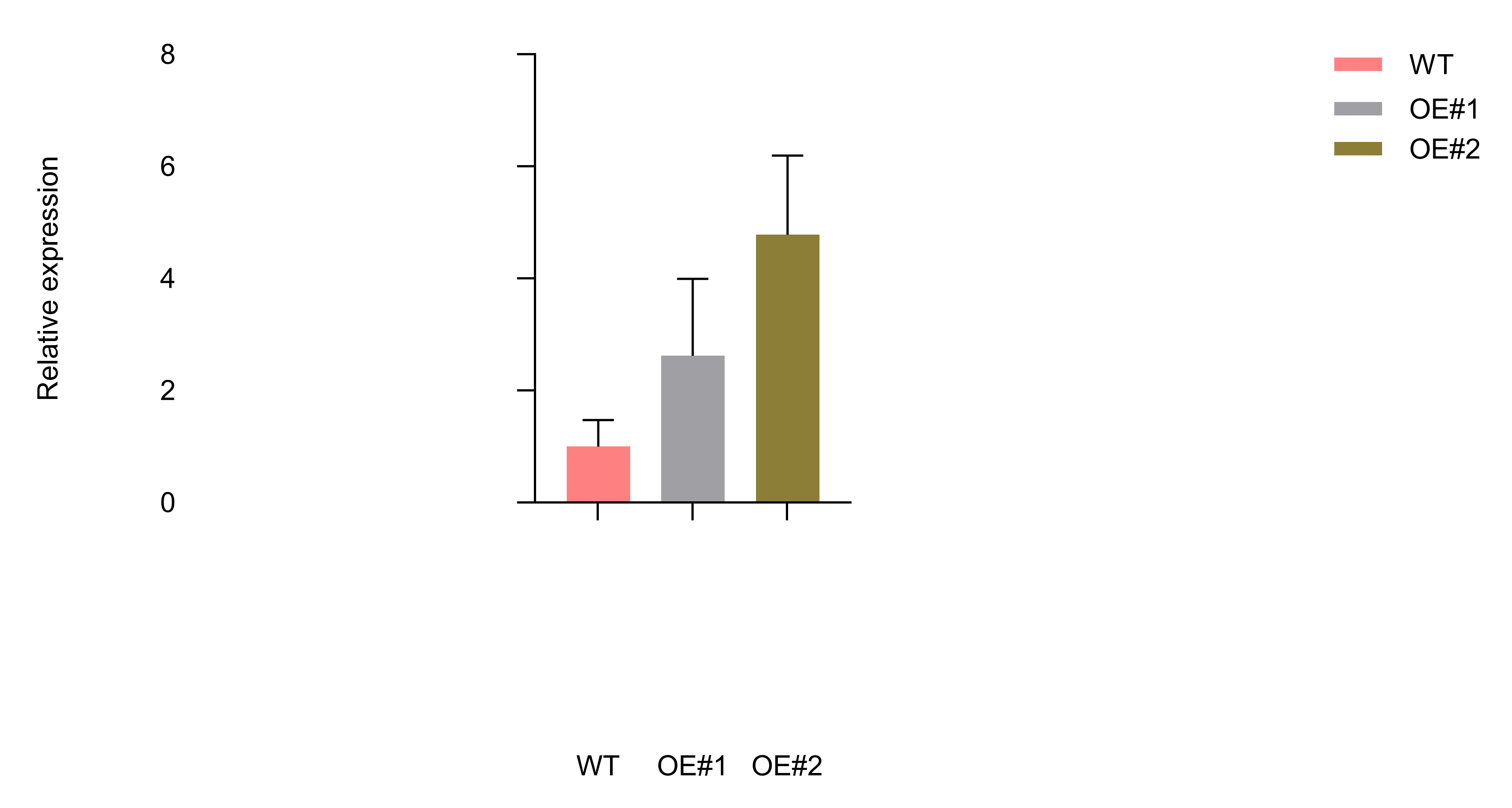

WT

*sec-4*

*sec-5*

WT

OE#1

OE#2

*sec-4*

*sec-5*

P

(a) Schematic genomic structure of the SEC locus. Exons are shown as purple boxes, introns are indicated as lines, UTR is shown as purple black box. T-DNA insertion positions are indicated by triangles. LB represents the left border primer of the T-DNA insertion. LP and RP represent the left and right genomic primers, respectively. P represents the SEC transcript detected by RT-qPCR.
(b) Identification of *sec* homozygotes. (M: Marker. The molecular weight is 5000 bp. The lane 1 and 2 in the left panel are *sec-4* mutants. The lane 2 in the right panel is the *sec-5* mutant).
(c) RT-qPCR analysis of SEC mRNA levels in *sec-4* and *sec-5* mutants. The expression level was normalized to that of *EF1-α*, a reference gene for RT-qPCR. Independent biological experiments were repeated three times. Data shown are means ± SD.
(d) RT-qPCR analysis of *SEC* mRNA levels in *SEC*-overexpression (OE) lines. The expression level was normalized to that of *EF1-α*, a reference gene for RT-qPCR. Independent biological experiments were repeated three times. Data shown are means ± SD.
