## Supplemental Tables for "The O-glycosyltransferase SECRET AGENT Participates in Abscisic Acid-Induced Microtubule Remodeling and Stomatal Closure in Arabidopsis thaliana"

List of primers used in this study:

Supplemental Table S1 Identification of *sec-4* (SALK_106339) mutants

| Primer name | Primer sequence (5′-3′) |
| --- | --- |
| LP | AAGGATCAGCTGTGAAGATGC |
| RP | TTGTATGGGGAGAGCATCAAG |
| LB | ATTTTGCCGATTTCGGAAC |

Supplemental Table S2 Identification of *sec-5* (SALK_034290) mutants

| Primer name | Primer sequence (5′-3′) |
| --- | --- |
| LP | TCATGAATCAATCCTTGAGCC |
| RP | TTTCGATGTCCCTTCTTTGTG |
| LB | ATTTTGCCGATTTCGGAAC |

Supplemental Table S3 RT-qPCR

| Primer name | Primer sequence (5′-3′) |
| --- | --- |
| *SEC*-F | GATCAGTGTTTGGGATGCAC |
| *SEC*-R | AGAAACGTCCAGGAAATGCT |
| *EF-1α*-F | TGAGCACGCTCTTCTTGCTTTCA |
| *EF-1α*-R | GGTGGTGGCATCCATCTTGTTACA |

Supplemental Table S4 Yeast two-hybrid assay

| Primer name | Primer sequence (5′-3′) |
| --- | --- |
| BD-*SEC*-F | AGGCCGAATTCCCGGGGATCCTGATCTCGTCCAAAAACGGA |
| BD-*SEC*-R | CTAGTTATGCGGCCGCTGCAGTCTGTCATGTGGGAATTCTA |
| *TUA4*-F | TCATAAACGCCCTTCGTCTTCTT |
| *TUA4*-R | ATTACCAGAAAGGCAGAAACGAT |
| AD-*TUA4*-F | GCCATGGAGGCCAGTGAATTCATGAGAGAGATCCTTCACATTC |
| AD-*TUA4*-R | CAGCTCGAGCTCGATGGATCCAGTCTCATAATCTCCCTCCTCT |

Supplemental Table S5 Bimolecular fluorescence complementation (BiFC) assay

| Primer name | Primer sequence (5′-3′) |
| --- | --- |
| *SEC*-nYFP-F | GAGAACACGGGGGACTCTAGAATGATCTCGTCCAAAAACGG |
| *SEC*-nYFP-R | GCCCTTGCTCACCATGGATCCTCTGTCATGTGGGAATTCTAGGT |
| *TUA4*-cYFP-F | GAGAACACGGGGGACTCTAGAATGAGAGAGATCCTTCACATTC |
| *TUA4*-cYFP-R | CTTCTGCTTGTCCATGGATCCAGTCTCATAATCTCCCTCCTCT |
